# Efficient sequence-specific isolation of DNA fragments and chromatin by *in vitro* enChIP technology using recombinant CRISPR ribonucleoproteins

**DOI:** 10.1101/033241

**Authors:** Toshitsugu Fujita, Miyuki Yuno, Hodaka Fujii

## Abstract

The clustered regularly interspaced short palindromic repeat (CRISPR) system is widely used for various biological applications, including genome editing. We developed engineered DNA-binding molecule-mediated chromatin immunoprecipitation (enChIP) using CRISPR to isolate target genomic regions from cells for their biochemical characterization. In this study, we developed *“in vitro* enChIP” using recombinant CRISPR ribonucleoproteins (RNPs) to isolate target genomic regions. *in vitro* enChIP has the great advantage over conventional enChIP of not requiring expression of CRISPR complexes in cells. We first demonstrate that *in vitro* enChIP using recombinant CRISPR RNPs can be used to isolate target DNA from mixtures of purified DNA in a sequence-specific manner. In addition, we show that this technology can be employed to efficiently isolate target genomic regions, while retaining their intracellular molecular interactions, with negligible contamination from irrelevant genomic regions. Thus, *in vitro* enChIP technology is of potential use for sequence-specific isolation of DNA, as well as for identification of molecules interacting with genomic regions of interest *in vivo* in combination with downstream analysis.

## Introduction

The advent of engineered DNA-binding molecules, such as zinc finger proteins, transcription activator-like effector (TAL or TALE) proteins, and the clustered regularly interspaced short palindromic repeat (CRISPR) system, has enabled high efficiency genome editing (Gaj et al., 2013; Mali et al., 2013; Sun & Zhao, 2013; Doudna & Charpentier, 2014; Harrison et al., 2014; Hsu et al., 2014; Pauwels et al., 2014; Wijshake et al., 2014). Moreover, the use of such engineered DNA-binding molecules is not limited to genome editing; they are used for various applications, including artificial transcriptional regulation, epigenetic modification, locus imaging, and isolation of specific genomic regions for the analysis of locus-specific genome functions (Mali et al., 2013; Harrison et al., 2014; Hsu et al., 2014; Fujii & Fujita, 2015; Fujita & Fujii, 2015a). Among these engineered DNA-binding molecules, the CRISPR system is the most convenient, economical, and time-efficient and has rapidly risen to prevail over technologies with similar functions since its introduction for use in genome editing.

To elucidate the molecular mechanisms underlying genome functions such as transcription and epigenetic regulation, it is necessary to identify molecules interacting with genome regions of interest *in vivo.* To this end, we developed locus-specific chromatin immunoprecipitation (locus-specific ChIP) technologies, consisting of insertional ChIP (iChIP) and engineered DNA-binding molecule-mediated ChIP (enChIP) (Hoshino & Fujii, 2009; Fujita & Fujii, 2011, 2012, 2013a, 2013b, 2014a, 2014b, 2015b; Fujita et al., 2013, 2015a, 2015b). In enChIP, an engineered DNA-binding molecule, such as a TAL protein or the CRISPR complex (guide RNA (gRNA) and dCas9, a nuclease-deficient mutant of Cas9), is expressed, often as an epitope tag-fused form, for locus tagging in the cell to be analyzed. The targeted locus can be isolated by affinity purification using an antibody (Ab) against the epitope tag or the CRISPR complex itself (Fig. S1). Previously, we successfully used enChIP combined with mass spectrometry and RNA sequencing (RNA-seq) for the non-biased identification of proteins and RNAs interacting with the *interferon regulatory factor-1 (IRF-1)* promoter region and telomere regions *in vivo* (Fujita et al., 2013; Fujita & Fujii, 2013b, 2014a; Fujita et al., 2015b).

In addition to the aforementioned conventional *in vivo* enChIP, we reported previously that using enChIP with recombinant engineered DNA-binding molecules, such as a TAL protein, it is feasible to isolate a genomic region of interest from a cell (Fujita & Fujii, 2014b). This “in *vitro* enChIP” system has the great advantage over conventional enChIP of not needing to express engineered DNA-binding molecules *in vivo.* However, in the *in vitro* enChIP assay using the TAL protein, we observed significant degradation of the recombinant TAL protein, and the DNA yields from the target locus were modest (Fujita & Fujii, 2014b). Development of *in vitro* enChIP technology that facilitates more efficient and specific isolation of target genomic regions would allow the application of this technology to the faster and simpler identification of molecules interacting with target genomic regions.

In this study, we developed an *in vitro* enChIP method using recombinant CRISPR ribonucleoproteins (RNPs) and demonstrated that it can be used to isolate target DNA from mixtures of purified DNA. In addition, the system was applicable to the efficient and specific isolation of target genomic regions from cells lysates. Thus, *in vitro* enChIP using recombinant CRISPR RNPs has potential uses as a sequence-specific DNA isolation tool and, in combination with downstream analysis, in identification of molecules interacting with genomic regions of interest *in vivo.*

## Results and Discussion

### Overview of in vitro enChIP using recombinant CRISPR RNPs

Our protocol for *in vitro* enChIP using recombinant CRISPR RNPs is as follows: (1) A recombinant 3xFLAG-dCas9-D protein, r3xFLAG-dCas9-D, consisting of a 3xFLAG-tag, a nuclear localization signal (NLS), dCas9, and a Dock-tag, is incubated with synthetic gRNA (e.g., a chimeric single guide RNA (sgRNA)) to form recombinant CRISPR RNPs. (2) The cell to be analyzed is crosslinked with formaldehyde, or another crosslinker, if necessary. The cell is lysed, and the chromatin is fragmented. (3) The target genomic region is captured *in vitro* by the CRISPR RNPs and affinity-purified with anti-FLAG Ab conjugated to carriers. After isolation, the chromatin components (DNA, RNA, proteins, and other molecules) interacting with the target genomic region can be identified by downstream analysis, such as next-generation sequencing (NGS), microarray, or mass spectrometry (Fig. 1).

**Fig. 1.**
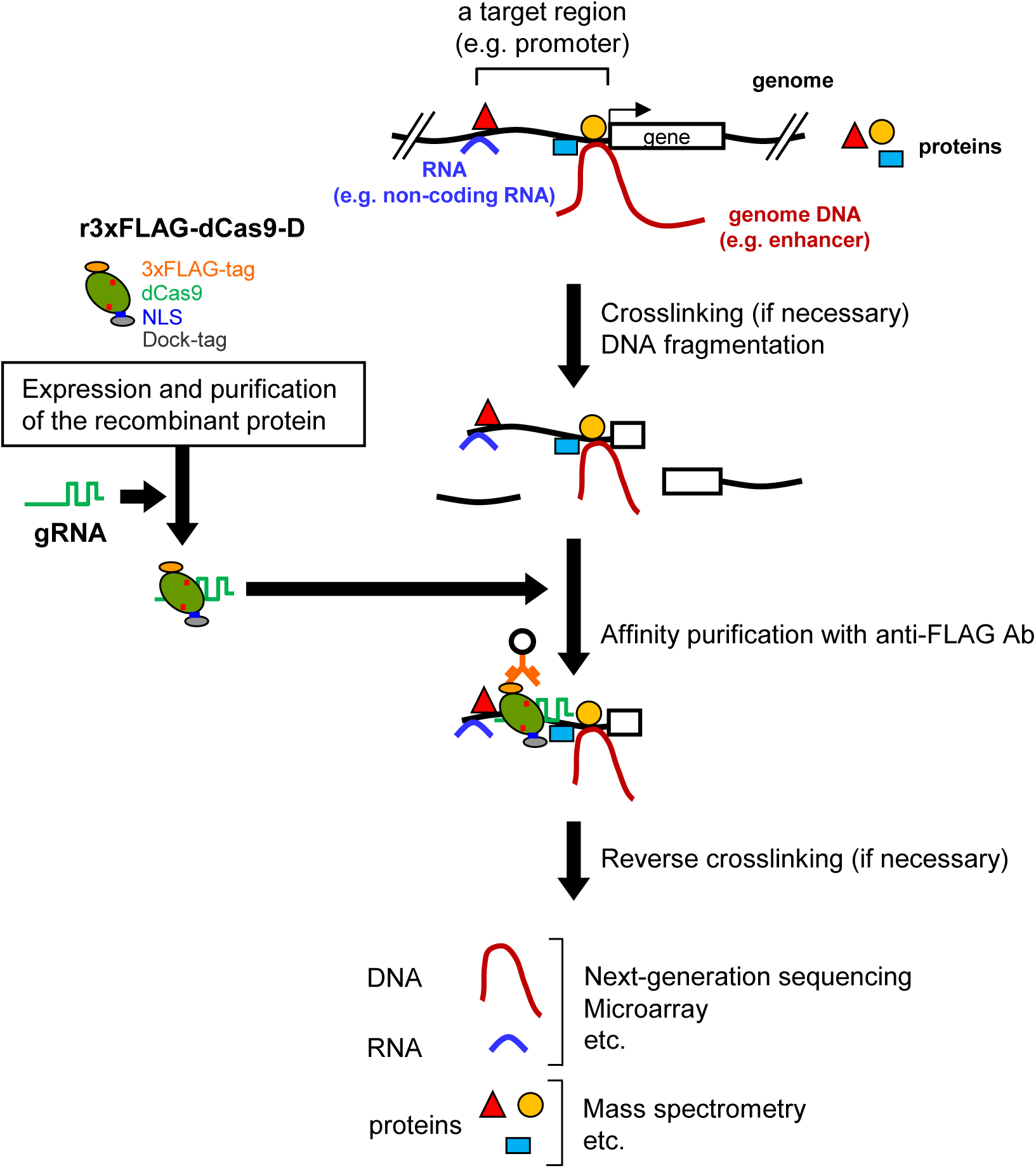
An overview of *in vitro* enChIP using recombinant CRISPR RNPs. Recombinant 3xFLAG-dCas9-D protein (r3xFLAG-dCas9-D) consisting of a 3xFLAG-tag epitope, a nuclear localization signal (NLS), dCas9, and a Dock-tag is incubated with gRNA to form CRISPR RNPs. The cell to be analyzed is crosslinked with formaldehyde or other crosslinkers, if necessary. The cell is lysed, and the genomic DNA is fragmented. The target genomic region is captured *in vitro* by the CRISPR RNPs and affinity-purified with anti-FLAG Ab, conjugated to carriers. Purification of the chromatin components (DNA, RNA, proteins, and other molecules) allows their identification by downstream analyses.

In addition to Ab-based purification, the biotin-avidin system (Diamandis & Christopoulos, 1991) can also be employed for affinity purification in *in vitro* enChIP, using r3xFLAG-dCas9-D and biotinylated gRNA (Fig. S2) (see below).

### Isolation of DNA fragments in a sequence-specific manner by in vitro enChIP using recombinant CRISPR RNPs

First, we generated the r3xFLAG-dCas9-D protein using the silkworm-baculovirus expression system. In contrast to the TAL protein (Fujita & Fujii, 2014b), r3xFLAG-dCas9-D could be purified without visible degradation (Fig. 2A). Next, we evaluated *in vitro* enChIP using the purified r3xFLAG-dCas9-D with synthesized gRNA targeting the *IRF-1* promoter region (Fig. 2B and C). Using fragmented purified 293T genomic DNA as input, the *IRF-1* region was enriched compared with non-target control regions (Fig. 2D and E). Approximately half of the total amount of the target region was efficiently enriched by *in vitro* enChIP combined with Ab-based (Fig. 2D) or biotin-avidin purification (Fig. 2E) system. In addition, we succeeded in specific isolation of a target PCR product from mixtures of DNA fragments (Fig. 2F). These results suggest that *in vitro* enChIP technology can be applied to sequence-specific concentration or removal of target DNA from mixtures of DNA such as fragmented genomic DNA and PCR products.

**Fig. 2.**
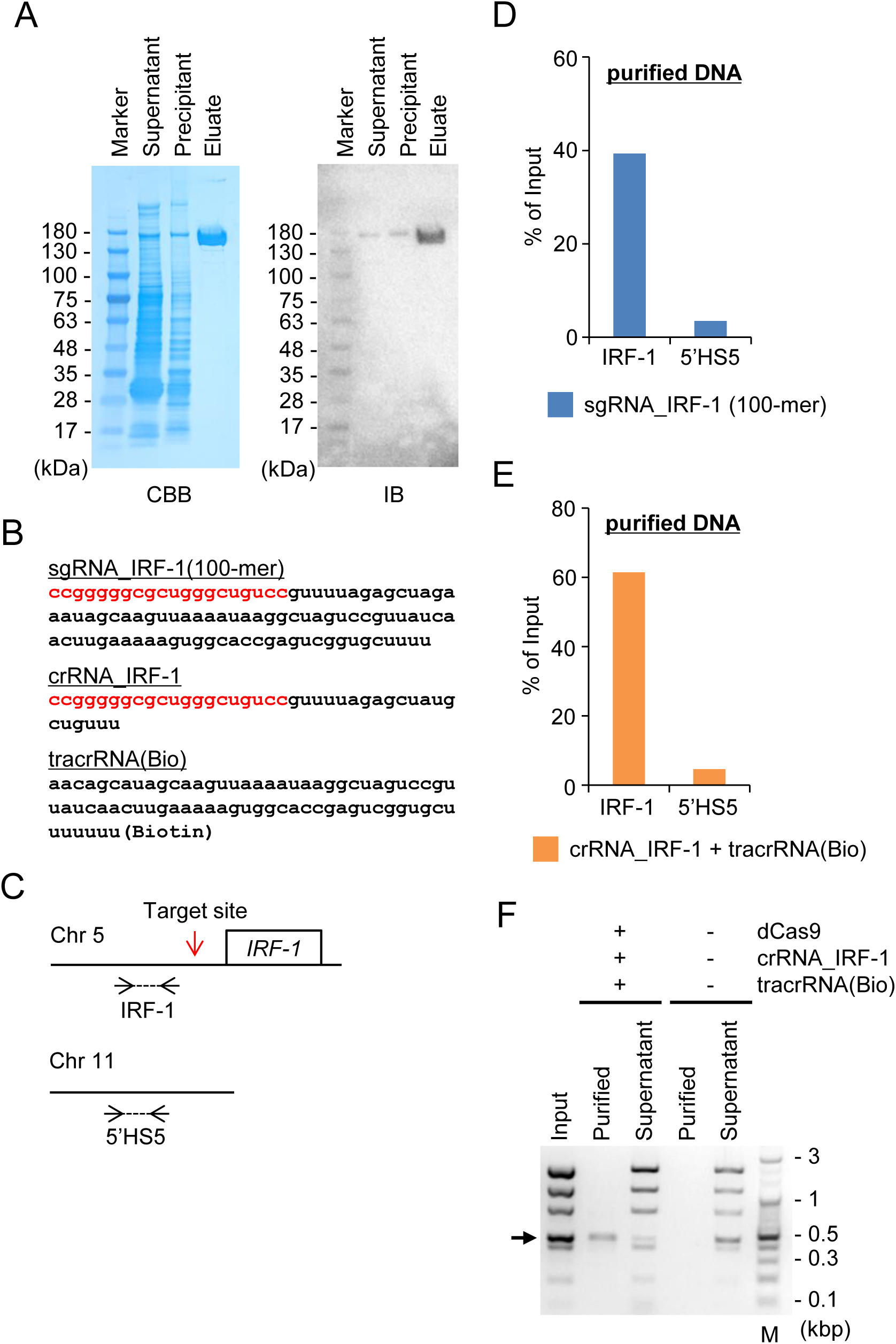
Isolation of DNA fragments in a sequence-specific manner by *in vitro* enChIP using recombinant CRISPR RNPs. **(A)** Preparation of r3xFLAG-dCas9-D. Purified proteins were subjected to SDS-PAGE and Coomassie Brilliant Blue staining (CBB) (left panel) or immunoblot analysis with anti-Dock Ab (IB) (right panel). Marker: molecular weight marker; Supernatant: the supernatant prepared from silkworm pupal homogenates; Precipitant: the insoluble precipitate prepared from silkworm pupal homogenates; Eluate: the eluate after affinity purification. **(B)** sgRNA, crRNA, and tracrRNA used in this study. Sequences highlighted in red are complementary to a target genomic sequence. **(C)** Positions of primers used in this study to amplify target (IRF-1, chromosome 5) and non-target control (5’HS5, chromosome 11) genome regions. The site targeted by *in vitro* enChIP is indicated by the red arrow. (**D** and **E**) The DNA yields from target (IRF-1) and non-target control (5’HS5) genome regions after *in vitro* enChIP combined with the Ab-based **(D)** or biotin-avidin **(E)** purification system. Purified genomic DNA fragments prepared from 293T cells were used for quantitative PCR. **(F)** Results of isolation of a target PCR product from a mixture of DNA fragments by *in vitro* enChIP combined with the biotin-avidin purification system (+) and a negative control where r3xFLAG-dCas9-D, crRNA, and tracrRNA were omitted (−). Input: the mixture of DNA fragments including the target PCR product (arrow); Purified: the DNA purified by *in vitro* enChIP; Supernatant: the supernatant after the *in vitro* enChIP procedure; M: molecular weight marker.

We envisage various applications of the *in vitro* enChIP technology as a sequence-specific DNA isolation tool. For example:

a. Sequence-specific purification of PCR amplicons and plasmids: In suboptimal experimental conditions, PCR may amplify not only a target DNA region but also non-specific sequences. If the molecular weight of the target amplicon is similar to those of non-specific products, it is difficult to separate the target amplicon from unwanted products by gel electrophoresis. In such cases, *in vitro* enChIP could be used to isolate the target amplicon in a sequence-specific manner. *in vitro* enChIP could also be used for sequence-specific isolation of plasmids containing target DNA from a cDNA or genomic DNA library.
b. Enrichment of DNA fragments with specific inserted elements: In addition, the technology could be applied for sequence-specific concentration of DNA fragments containing specific inserted elements, such as viral, plasmid, or transposon sequences, from purified genomic DNA; NGS analysis using DNA concentrated by *in vitro* enChIP could identify the genomic positions of the integrated sequences with fewer reads.
c. Removal of unwanted DNA in transcriptome analysis, metagenomics, and analysis of microbial flora: In transcriptome analysis using RNA-seq, cDNA synthesized from abundant ribosomal RNA may dominate sequencing reads. Similarly, in *de novo* genome sequencing, repetitive sequences such as telomeric repeats may complicate sequence assembly. In these cases, removal of such undesirable DNA by *in vitro* enChIP could improve the sequencing resolution obtained. For analysis of microbial flora, sequencing of microbial 16S rDNA is used for species identification. It can be difficult to accurately detect rare microbes by NGS because the 16S rDNA of major microbial populations dominates in the analysis. Removal of such dominant 16S rDNA by *in vitro* enChIP could facilitate identification of rare microbe species.

Currently, a DNA hybridization using biotinylated DNA probes combined with the biotin-avidin purification system is used for a number of the aforementioned applications (Diamandis & Christopoulos, 1991; Ito et al., 1992; Mangiapan et al., 1996; St. John & Quinn, 2008; Camara Teixeira et al., 2013; Gagan & Van Allen, 2015); however, *in vitro* enChIP technology is a potentially useful alternative method.

### Isolation of target chromatin complexes in a sequence-specific manner by in vitro enChIP using recombinant CRISPR RNPs

Finally, we applied the *in vitro* enChIP technology combined with the Ab-based purification system to the isolation of target genomic regions retaining molecular interactions from cells. When we used native chromatin prepared from the human 293T cell line and a 100-mer sgRNA, the *IRF-1* region was efficiently enriched compared with non-target control regions (Fig. 3A). The DNA yields (% of input) of the target region were approximately 0.5%. A shorter sgRNA, lacking 38 nucleotides at the 3’ end compared with the 100-mer sgRNA, was not effective in our experimental system (Fig. S3). Specific enrichment of the target region was also observed using crosslinked chromatin as input (Fig. 3B); the DNA yields of the crosslinked target region were comparable using sgRNA (100-mer) and the CRISPR RNA (crRNA)/trans activating crRNA (tracrRNA) complex for specific targeting (Figs. S3 and S4). Thus, these results clearly showed that it is feasible to isolate target chromatin complexes from cells in a sequence-specific manner using *in vitro* enChIP with recombinant CRISPR RNPs. Moreover, *in vitro* enChIP combined with the biotin-avidin purification system was also successfully applied for this purpose (Fig. 3C). Although the DNA yields of the target region were comparable between both purification systems, the Ab-based purification system showed more than 10× less contamination with non-target control regions (Fig. 3B and C). The elution step of the Ab-based purification system, in which the recombinant CRISPR RNPs are specifically eluted from Ab-coated carriers using 3xFLAG peptides, may contribute to the lower backgrounds achieved using this method (see Experimental procedures).

**Fig. 3.**
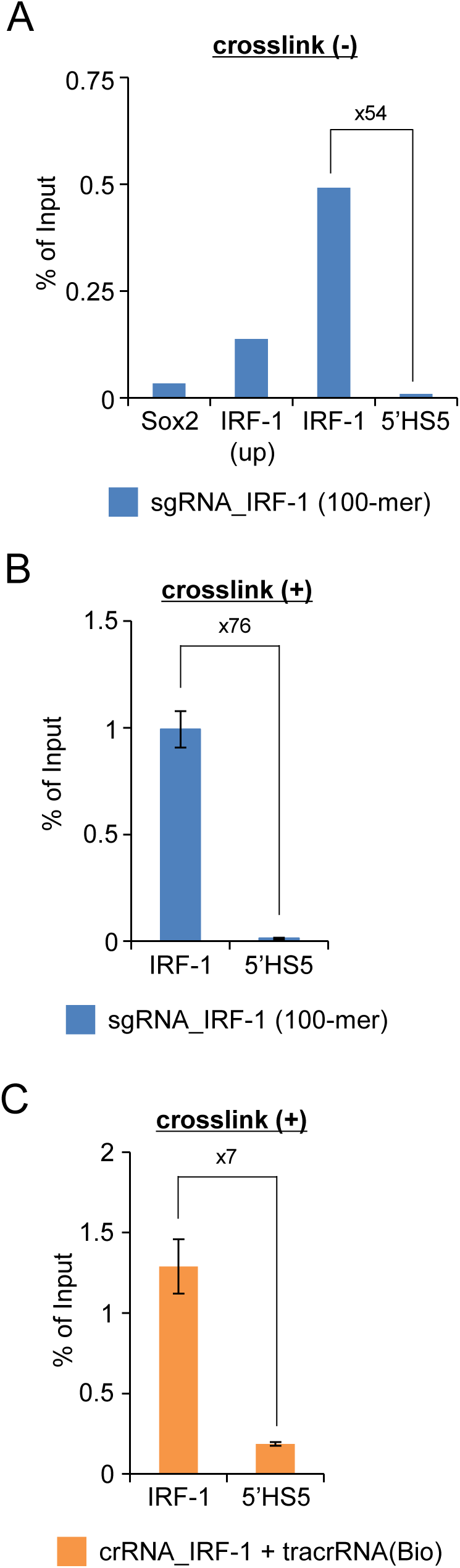
*In vitro* enChIP for isolation of target chromatin complexes in a sequence-specific manner. **(A)** Plot of DNA yields of *in vitro* enChIP with native chromatin prepared from 293T cells determined by quantitative PCR from target (IRF-1) and non-target (Sox2 and 5’HS5) genome regions. (**B** and **C**) DNA yields measured by quantitative PCR from samples subjected to *in vitro* enChIP with crosslinked chromatin prepared from 293T cells. The Ab-based purification system (**A** and **B**) or the biotin-avidin purification system (**C**) was employed. IRF-1: target genomic region; IRF-1 (up): 0.1 kbp upstream from the primer position for IRF-1; 5’HS5 and Sox2: non-target control genomic regions. The error bar represents the SEM of three *in vitro* enChIP experiments (**B** and **C**).

In contrast to *in vitro* enChIP using a recombinant TAL protein, where the DNA yields were only 0.01% of input (Fujita & Fujii, 2014b), *in vitro* enChIP using the recombinant CRISPR RNPs showed much higher DNA yields (~1%) of the target locus using both native and crosslinked chromatin input material (Fig. 3). Thus, *in vitro* enChIP using recombinant CRISPR RNPs could potentially be applied for the identification of molecules interacting with genomic regions of interest *in vivo.* Conventional enChIP, using CRISPR targeting the same site in the *IRF-1* promoter region, gave target DNA yields of ~10% (Fujita & Fujii, 2013b, 2014a). Since conventional enChIP can identify proteins interacting with the promoter region using ~ 1 × 10^8^ cells (Fujita & Fujii, 2013b, 2014a), it may be possible to identify proteins interacting with the region by *in vitro* enChIP using recombinant CRISPR RNPs from ~ 1 × 10^9^ cells. Smaller numbers of cells would be sufficient for analysis of RNA and DNA interacting with a target genomic region.

### Conclusions

In this study, we developed *in vitro* enChIP technology using recombinant CRISPR RNPs. *In vitro* enChIP does not require expression of the engineered DNA-binding molecules *in vivo.* This is a great advantage over conventional enChIP technology, since *in vitro* enChIP can be employed without the time-consuming steps required to generate cells expressing tagged DNA-binding molecules or the preparation of transgenic organisms, such as mice. *in vitro* enChIP might also be useful in situations where the generation of transgenic organisms is difficult or impossible, for example, when non-model organisms or pathogens are under study. Moreover, it is not necessary to consider undesirable side-effects such as CRISPR interference (CRISPRi) (Bikard et al., 2013; Qi et al., 2013; Zhao et al., 2014). Our results indicate that *in vitro* enChIP using recombinant CRISPR RNPs is a potentially useful sequence-specific DNA isolation tool in biochemistry and molecular biology, applicable to the identification of molecules interacting with specific genomic regions. Using *in vitro* enChIP technology, we are now performing experiments to identify molecules interacting with genomic regions of interest *in vivo* to elucidate the molecular mechanisms underlying genome functions.

## Experimental procedures

### Preparation of r3xFLAG-dCas9 and gRNA

Preparation of r3xFLAG-dCas9 and gRNA are shown in Supplemental Experimental procedures.

### in vitro enChIP using r3xFLAG-dCas9-D/gRNA RNPs

Two micrograms of r3xFLAG-dCas9-D were incubated with 2 μg of anti-FLAG M2 Ab (Sigma-Aldrich) conjugated to 30 μl of Dynabeads-Protein G (Thermo Fisher Scientific) in PBS-T-BSA (PBS, 0.05% Tween-20, and 0.1% BSA) at 4°C for 2–3 h. After washing with PBS-T (PBS and 0.05% Tween-20), the Dynabeads were incubated with the annealed sgRNA or crRNA/tracrRNA at 37°C for 10 min in 100 μl of *in vitro* CRISPR buffer (20 mM HEPES pH 7.5, 150 mM KCl, 0.1 mM EDTA, 10 mM MgCl_2_, and 0.5 mM DTT) and used for *in vitro* enChIP.

Purified 293T genomic DNA (5 μg) was sonicated in 800 μl of *in vitro* CRISPR buffer as reported previously (Fujita & Fujii, 2013b). The average length of DNA fragments was approximately 1 kb. The sonicated genomic DNA was pre-cleared with 2 μg of normal mouse IgG (Santa Cruz Biotechnology) conjugated to 30 μl of Dynabeads-Protein G. The supernatant was incubated with the prepared r3xFLAG-dCas9-D/gRNA RNP complex conjugated to Dynabeads and 40 U/ml RNasin plus RNase inhibitor (Promega) at 37°C for 20 min. The Dynabeads were washed as described previously (Fujita & Fujii, 2014b) with some modifications. The captured chromatin complexes were eluted with 60 μl of elution buffer (500 μg/ml 3xFLAG peptide (Sigma-Aldrich), 50 mM Tris pH 7.5, 150 mM NaCl, and 0.1% IGEPAL-CA630) at 37°C for 20 min. The eluate was incubated with 2 μl of 10 μg/μl RNase A at 37°C for 1 h and then with 5 μl each of 10% SDS and 20 mg/ml Proteinase K (Roche) at 45°C for 2 h. DNA was purified using ChIP DNA Clean & Concentrator (Zymo Research) and analyzed by real-time PCR as described previously (Fujita & Fujii, 2013b). The primers used in this study are shown in Table S1.

For *in vitro* enChIP with native chromatin, native chromatin digested with micrococcal nuclease was prepared as reported previously (Fujita & Fujii, 2014b). The native chromatin prepared from 4 × 10^6^ 293T cells (300 μl) was mixed with 1.1 ml of *in vitro* CRISPR buffer including cOmplete protease inhibitor cocktail (EDTA-free) and used for *in vitro* enChIP as described above.

For *in vitro* enChIP with crosslinked chromatin, chromatin was crosslinked and fragmented by sonication as reported previously (Fujita & Fujii, 2013b), except that *in vitro* CRISPR buffer was used in the sonication step. Fragmented crosslinked chromatin prepared from 4 × 10^6^ 293T cells (160 μl) was mixed with 340 μl of *in vitro* CRISPR buffer including cOmplete protease inhibitor cocktail (EDTA-free) and used for *in vitro* enChIP as described previously (Fujita & Fujii, 2014b) with some modifications and with the inclusion of a reverse crosslinking step.

### in vitro enChIP using a r3xFLAG-dCas9-D/biotinylated gRNA RNP

Instead of non-labeled tracrRNA, tracrRNA biotinylated at the 3’ position was used. r3xFLAG-dCas9-D was incubated with a crRNA and biotinylated tracrRNA complex at 37°C for 5 min in 44.5 μl of *in vitro* CRISPR buffer and used for *in vitro* enChIP.

Sonicated genomic DNA (2 μg) purified from 293T cells in 500 μl of *in vitro* CRISPR buffer was pre-cleared with 30 μl of Dynabeads-streptavidin T1 (Thermo Fisher Scientific). The supernatant was incubated with the r3xFLAG-dCas9-D/biotinylated gRNA RNP complex at 37°C for 20 min and then mixed with 30 μl of Dynabeads-streptavidin T1 at 4°C for 1 h. The Dynabeads were washed as described previously (Fujita & Fujii, 2014b) with some modifications. The captured chromatin complexes were treated with RNase A and Proteinase K. DNA was purified using ChIP DNA Clean & Concentrator.

For the isolation of a PCR product, the *IRF-1* promoter region, including the target site for gRNA, was amplified by PCR and purified. The primers used for PCR amplification are shown in Table S1. PCR product (0.1 μg) and Low DNA Mass Ladder (1.2 μg; Thermo Fisher Scientific) were used for *in vitro* enChIP as described above. Before and after the *in vitro* enChIP procedure, DNA was analyzed by 2% agarose gel electrophoresis.

For *in vitro* enChIP using crosslinked chromatin, crosslinked and sonicated chromatin prepared from 4 × 10^6^ 293T cells (160 μl) was mixed with 340 μl of *in vitro* CRISPR buffer including cOmplete protease inhibitor cocktail (EDTA-free) and used for *in vitro* enChIP as described previously (Fujita & Fujii, 2014b) with some modifications and with the inclusion of a reverse crosslinking step.

## Acknowledgements

This work was supported by the Takeda Science Foundation (T.F.); the Asahi Glass Foundation (H.F.); the Kurata Memorial Hitachi Science and Technology Foundation (T.F. and H.F.); a Grant-in-Aid for Scientific Research (C) (#15K06895) (T.F.); a Grant-in-Aid for Scientific Research on Innovative Areas “Transcription Cycle” (#25118512 & #15H01354) (H.F.); a Grant-in-Aid for Scientific Research (B) (#15H04329) (H.F. and T.F.); and a Grant-in-Aid for Exploratory Research (#26650059) (H.F.) from the Ministry of Education, Culture, Sports, Science and Technology of Japan.

## Conflicts of interest

T.F. and H.F. have filed a patent application relating to enChIP (Patent name: “Method for isolating specific genomic regions using DNA-binding molecules recognizing endogenous DNA sequences”; Patent number: WO2014/125668).

## Supplemental Materials

## Inventory

Supplemental Experimental Procedures

Supplemental Figure Legends

Supplemental Figures S1 - S4

Supplemental Table S1

## Supplemental Experimental Procedures

### Preparation of r3xFLAG-dCas9-D

r3xFLAG-dCas9-D was expressed using the silkworm-baculovirus expression system (ProCube^®^, http://procube.sysmex.co.jp/eng/) (Sysmex Corporation) as described previously (Fujita & Fujii, 2014b). Briefly, the coding sequence of the 3xFLAG-dCas9 from the 3xFLAG-dCas9/pCMV7.1 plasmid (Addgene #47948) was inserted into the transfer vector pM31a (Sysmex Corporation) to fuse with the Dock-tag at its C-terminus (r3xFLAG-dCas9-D). As reported previously (Fujita & Fujii, 2014b), the resultant plasmid was used for expression of r3xFLAG-dCas9-D in a silkworm pupa, and the expressed protein was affinity-purified.

### Preparation of gRNA complexes

sgRNAs and tracrRNAs were synthesized by Gene Design, Inc. crRNA was synthesized by FASMAC. The synthesized RNAs were diluted with nuclease-free water to a final concentration of 10 μM. Two microliters of sgRNA or 2 μl each of crRNA and tracrRNA were incubated at 100°C for 2 min and cooled down to room temperature.

## Supplemental Figure Legends

**Fig. S1.**
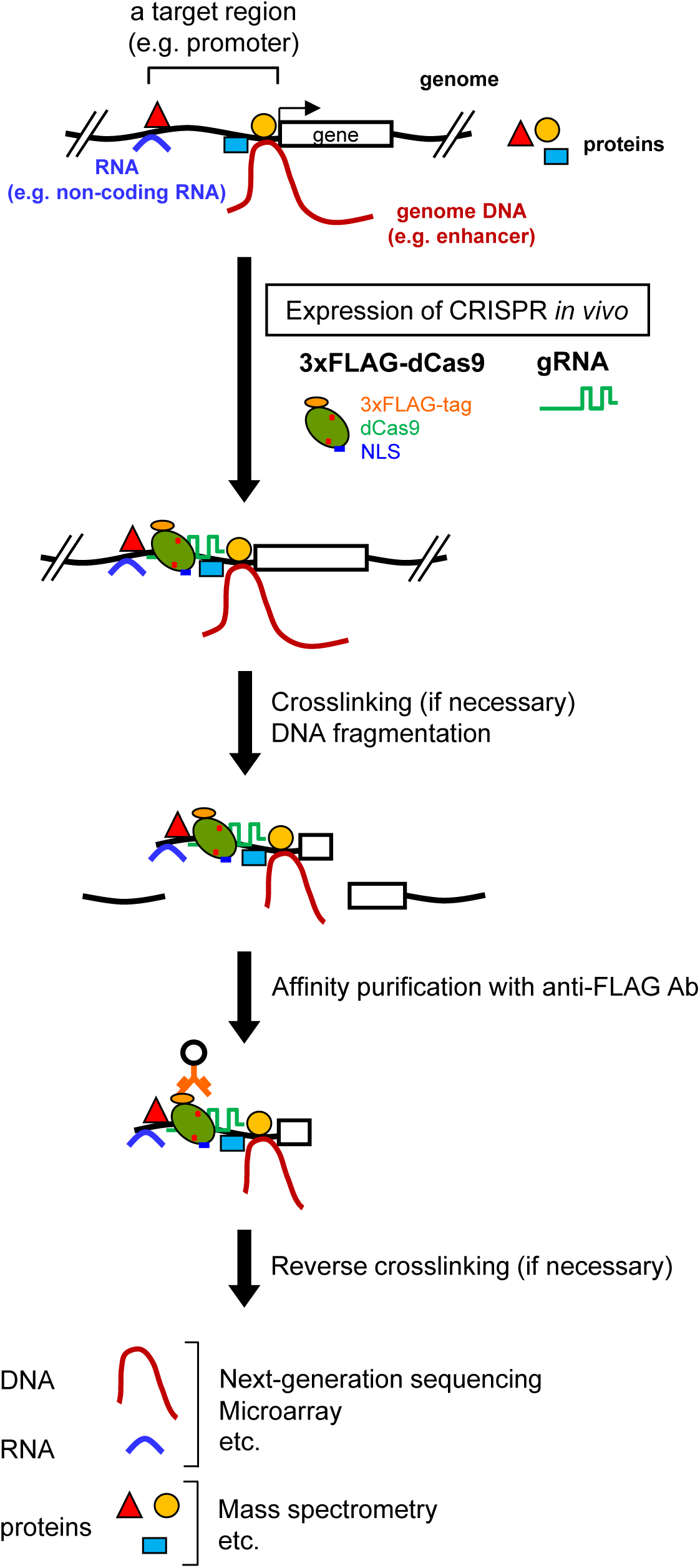
A protocol for conventional enChIP using CRISPR. 3xFLAG-dCas9, consisting of a 3xFLAG-tag epitope, a nuclear localization signal (NLS), and dCas9, is expressed together with gRNA (e.g., sgRNA) in the cell to be analyzed. The cell is crosslinked with formaldehyde or other crosslinkers, if necessary. The cell is lysed, and the genomic DNA is fragmented. The genomic region targeted by the CRISPR complex is affinity-purified with anti-FLAG Ab conjugated to carriers. Purification of the chromatin components (DNA, RNA, proteins, and other molecules) allows their identification by downstream analyses.

**Fig. S2.**
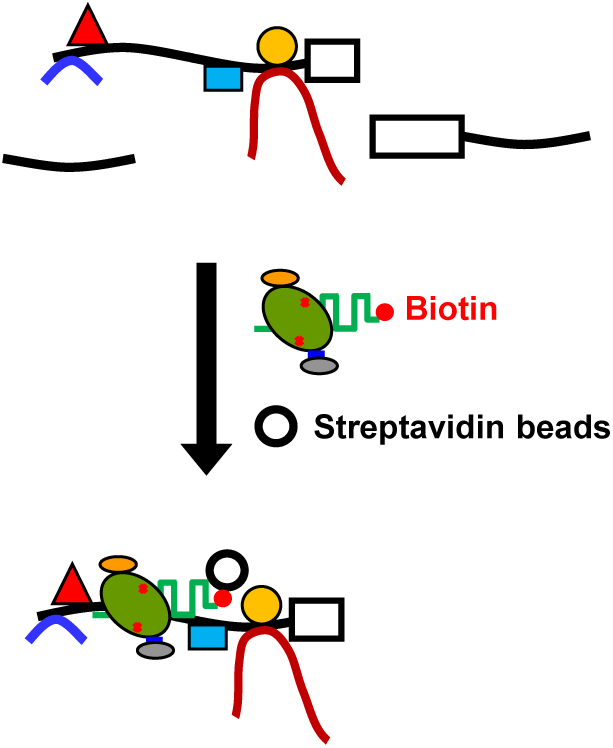
A protocol for *in vitro* enChIP with biotinylated gRNA combined with the biotin-avidin purification system.

**Fig. S3.**
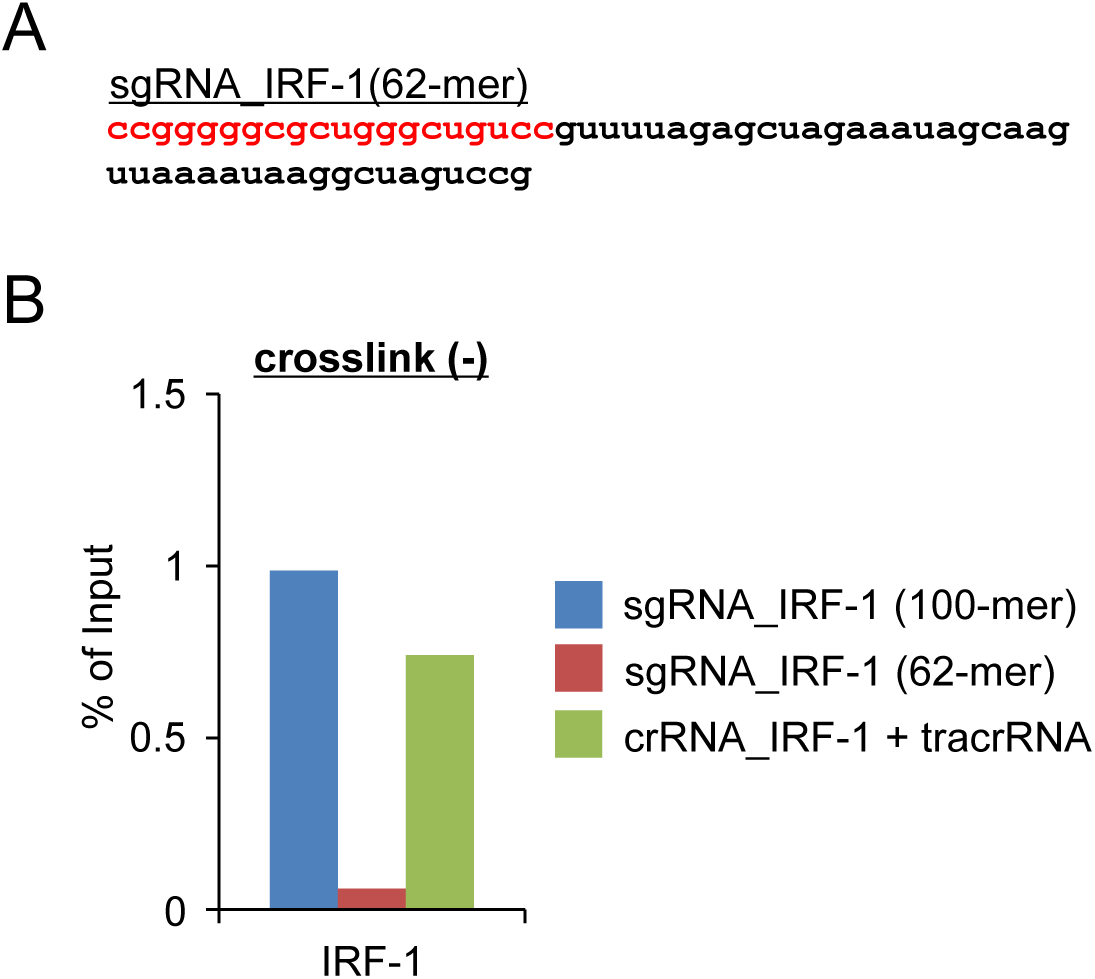
*in vitro* enChIP with sgRNA (62-mer). (**A**) sgRNA (62-mer) used in this study (sgRNA and crRNA used in this study are shown in Figure 2B). Sequences highlighted in red are complementary to a target genomic sequence. (**B**) DNA yields of *in vitro* enChIP with native chromatin prepared from 293T cells.

**Fig. S4.**
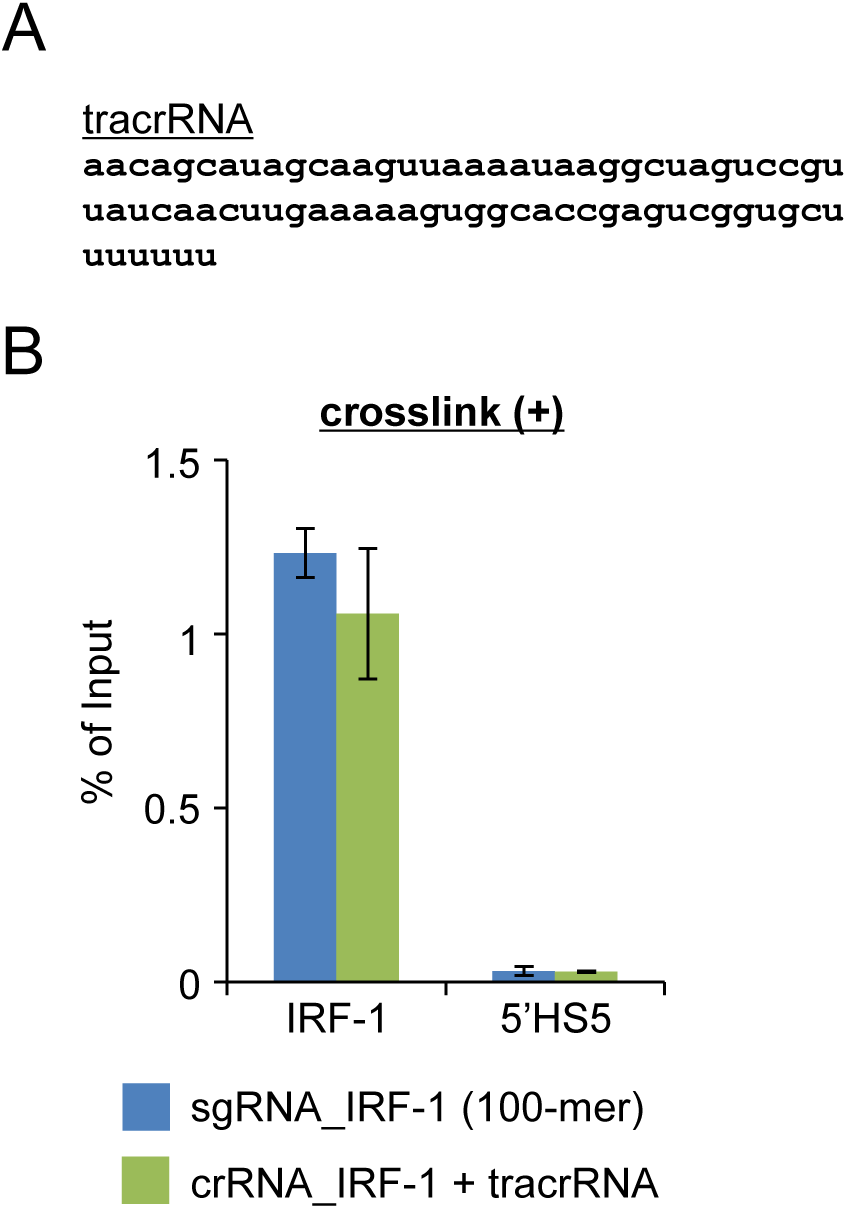
*in vitro* enChIP using recombinant CRISPR RNPs. (**A**) tracrRNA used in this study (sgRNA and crRNA used in this study are shown in Figure 2B). (**B**) DNA yields of *in vitro* enChIP with crosslinked chromatin prepared from 293T cells. The error bar represents the range of yields from duplicate *in vitro* enChIP experiments.

**Table S1.**
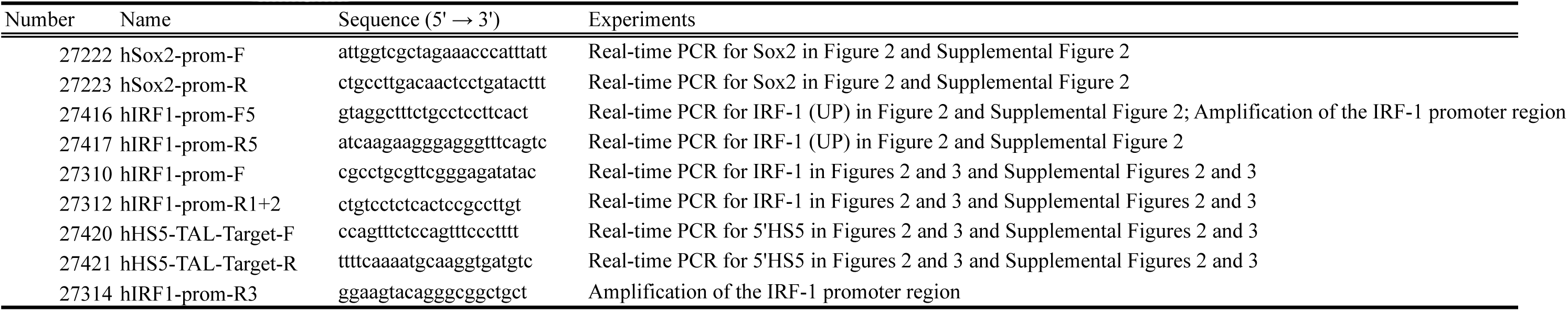
Primers used in this study

